# In Silico Screening of Neem-Derived Phytochemicals Targeting the Avr1 (SIX4) Effector Protein of *Fusarium oxysporum* for Fusarium Wilt Management

**DOI:** 10.64898/2025.12.25.696551

**Authors:** Raihana Sultana, Rofiqul Islam Nayem, Md. Mahidul Islam Masum

**Affiliations:** Department of Plant Pathology, Gazipur Agricultural University, Gazipur 1706, Bangladesh

**Author notes:** Corresponding author’s. **Author Contributions Mst Raihana Sultana:** Conducted the molecular docking experiments, performed ADME and toxicity analyses, analyzed and interpreted the computational results, and contributed to manuscript revision; **Rofiqul Islam Nayem:** Assisted in ligand selection and preparation, validated docking results, curated data, prepared figures and tables, and contributed to manuscript writing and revision; **Dr. Md. Mahidul Islam Masum:** Conceived and supervised the study, provided methodological and scientific guidance, critically reviewed and edited the manuscript, and approved the final version for submission.

**Keywords:** Fusarium wilt, Avr1 effector protein, molecular docking, neem phytochemicals, ADME, toxicity prediction

## Abstract

Fusarium wilt, caused by *Fusarium oxysporum*, is a destructive soil-borne disease that severely limits crop productivity worldwide. It’s effector protein Avr1 plays a key role in suppressing host immune responses. In this study, an integrated *in silico* approach was employed to identify potential natural inhibitors of Avr1 using neem (*Azadirachta indica*) derived phytochemicals. Ten compounds were screened through molecular docking, ADME profiling, and toxicity prediction. Docking results revealed binding affinities ranging from −6.3 to −5.5 kcal/mol, with nimbiol showing the strongest interaction (−6.3 kcal/mol). The highest-ranking protein–ligand complex was stabilized by hydrophobic and electrostatic interactions involving key residues such as ARG180, VAL158, LYS182, and TYR134, indicating a favorable binding environment. ADME analysis demonstrated acceptable pharmacokinetic properties and drug-likeness for several compounds, specially nimbiol and gedunin, with high gastrointestinal absorption and compliance with Lipinski’s rule of five. Toxicity prediction indicated moderate hepatotoxicity and low carcinogenic potential, although elevated immunotoxicity was observed across most ligands. Overall, the findings align with existing studies on neem’s antifungal potential and highlight nimbiol and gedunin as promising lead compounds for further experimental validation against Fusarium wilt.

## 1.0 Introduction

Fusarium wilt is one of the most destructive plant diseases affecting a wide range of economically important crops worldwide (Knights & Hobson, 2016), including banana, watermelon, cotton, tomato, and several horticultural and forest species. The disease is caused by the soil-borne fungal pathogen *Fusarium oxysporum*, which is characterized by its host-specific formae speciales (*Fusarium Wilt* | *Greenlife Crop Protection Africa*, 2022). It also has the ability to persist in soil for extended periods through durable survival structures (Gordon, 2017). Recent reports from different geographical regions have highlighted the increasing incidence and expanding host range of Fusarium wilt, resulting in significant yield losses and posing a serious threat to sustainable agricultural production (Nguyen et al., 2025; Thayne Munhoz et al., 2024; Raman Thangavelu et al., 2024; Sampaio et al., 2020).

Conventional management strategies for Fusarium wilt rely heavily on chemical fungicides and resistant cultivars (Khan et al., 2017). However, the effectiveness of these approaches is often limited by the emergence of fungicide resistance, environmental concerns along with the pathogen’s genetic variability. These challenges necessitate the exploration of alternative, environmentally friendly strategies that target key molecular components involved in pathogen virulence and host colonization.

The pathogenic success of *F. oxysporum* is largely attributed (Mwangi et al., 2020) to its ability to secrete a diverse set of effector proteins. These proteins facilitate the host invasion and suppress the plant immune responses. These proteins, commonly referred to as SIX (Secreted In Xylem) effectors (Fleur Gawehns et al., 2015), are delivered into the host vascular system during infection and play critical roles in disease development. Among them, Avr1, also known as SIX4, has been identified as an important virulence-associated effector protein in *F. oxysporum* f. sp. *lycopersici* and related formae speciales (TAKKEN & REP, 2010).

Avr1 is known to interfere with host defense signaling pathways and contribute to successful colonization of host tissues (Rafiqi et al., 2010). Structural and functional studies have suggested that Avr1 participates in molecular interactions essential for pathogenicity, making it an attractive target for antifungal intervention (Houterman et al., 2008). Targeting effector proteins such as Avr1 represents a promising strategy to attenuate pathogen virulence without directly exerting lethal pressure on the pathogen (DE WIT et al., 2009), thereby potentially reducing the risk of resistance development.

Natural products derived from medicinal plants have long been recognized as valuable sources of bioactive compounds for plant disease management. *Azadirachta indica* (neem) is particularly notable for its broad-spectrum antimicrobial, antifungal, and insecticidal properties (A. Akpuaka et al., 2013). Neem-derived phytochemicals, including limonoids (Abdelgaleil et al., 2005) and flavonoids such as nimbiol, gedunin, nimbin, and related compounds. These exhibit diverse chemical structures and biological activities that make them promising candidates for antifungal drug discovery (Janja, 2025).

Previous studies have reported the antifungal potential of neem extracts and individual compounds against various plant pathogens (Fernandes et al., 2019). However, the molecular mechanisms underlying their interaction with specific fungal virulence factors remain poorly understood. (Bhattacharyya & Jha, 2011; Fernandes et al., 2019). In particular, the potential of neem-derived compounds to target effector proteins of *F. oxysporum*, such as Avr1, has not been systematically explored.

Given the central role of Avr1 in Fusarium wilt pathogenesis and the known bioactivity of neem-derived phytochemicals, this study aims to investigate the interaction between selected neem compounds and the Avr1 effector protein using an in silico approach. Molecular docking was employed to screen a curated library of phytochemicals against Avr1 to identify potential high-affinity binders. The top-ranked compounds were further evaluated for their pharmacokinetic properties through ADME analysis and assessed for potential toxicity risks using predictive computational tools. Additionally, detailed protein–ligand interaction analyses were performed to elucidate the binding modes and key amino acid residues involved in complex stabilization.

This integrative computational study provides insights into the potential of neem-derived phytochemicals as lead compounds targeting the Avr1 effector protein. Ultimately, lays the foundation for future experimental validation and development of eco-friendly strategies for Fusarium wilt management.

## 2.0 Materials and Methods

### 2.1 Ligand Selection

A total of ten phytochemicals were selected from *Azadirachta indica* (neem) based on documented antifungal activity and structural diversity. These compounds included nimbiol, 6-deacetylnimbin, kulinone, methyl kulonate, kulactone, gedunin, kulolactone, 6beta-hydroxystigmast-4-en-3-one, methyl 2,5-dihydroxycinnamate, and sugiol. Canonical SMILES and PubChem CID identifiers were retrieved for each compound to ensure correct structural representation.

### 2.2 Target Protein Preparation

The effector protein Avr1 (SIX4) from *Fusarium oxysporum* f. sp. *lycopersici* was selected as the molecular target due to its critical role in virulence. The three-dimensional structure of Avr1 was obtained from PDB and prepared for docking by removing water molecules. Polar hydrogens were also added and protonation states at physiological pH using AutoDockTools were also optimized. Energy minimization was performed to relieve any steric clashes prior to docking.

### 2.3 Molecular Docking

Protein–ligand docking was carried out using AutoDock Vina. The binding site was defined based on known active residues identified from structural and literature studies. For each ligand, multiple docking poses were generated. And then the top-ranked pose was selected based on lowest binding free energy (kcal/mol) and RMSD values. RMSD/ub (upper bound) and RMSD/ib (internal bound) were calculated to ensure pose reliability. Docking results were visualized using PyMOL and Discovery Studio Visualizer.

### 2.4 ADME Analysis

Pharmacokinetic properties of the top ten ligands were predicted using SwissADME. Parameters analyzed included molecular weight, number of hydrogen bond acceptors and donors, lipophilicity (iLOGP), water solubility (Log S), gastrointestinal (GI) absorption, blood–brain barrier (BBB) permeability, Lipinski’s rule of five compliances, and PAINS (pan-assay interference compounds) alerts. These analyses helped identify compounds with favorable drug-likeness and bioavailability for further consideration.

### 2.5 Toxicity Prediction

Toxicity profiles of the selected ligands were evaluated using ProTox-II, which predicts hepatotoxicity, carcinogenicity, immunotoxicity, mutagenicity, and cytotoxicity based on machine-learning models. Compounds were assigned numerical toxicity scores, and active toxic endpoints were annotated for further interpretation.

### 2.6 Protein–Ligand Interaction Analysis

The highest-ranking Avr1–ligand complex was subjected to detailed interaction analysis using Discovery Studio Visualizer. Key contacts, including hydrogen bonds, hydrophobic interactions, electrostatic interactions, Pi–alkyl, and Pi–cation interactions, were identified and their distances measured. This analysis allowed a comprehensive understanding of the molecular interactions stabilizing the complex.

### 2.7 Data Representation

Tables were constructed to summarize docking affinities, ADME properties, toxicity predictions, and protein–ligand interactions. Figures of the top-ranked complexes were generated to illustrate binding orientations and interaction networks. All analyses were performed using default software parameters unless otherwise specified.

## 3.0 Results

### 3.1 Molecular Docking Analysis

Molecular docking was performed to evaluate the binding potential of ten neem-derived phytochemicals against the Avr1 (SIX4) effector protein of *Fusarium oxysporum*. The docking results revealed a range of binding affinities from –5.5 to –6.3 kcal/mol, with nimbiol exhibiting the highest binding affinity (–6.3 kcal/mol), followed closely by gedunin (–6.0 kcal/mol) (Table 2). These results indicate that these compounds can potentially form stable interactions within the predicted binding site of Avr1, suggesting inhibitory potential against the effector protein. The RMSD values for the docked complexes were within acceptable limits, confirming the reliability of the docking poses.

**Table 1.** Selected target disease, protein, and phytochemical sources.

| Target Disease | Pathogen | Target Protein | Protein Function | Natural Source | Compound |
| --- | --- | --- | --- | --- | --- |
| Fusarium wilt | *Fusarium oxysporum* | Avr1 (SIX effector) | Suppression of host immune response | *Azadirachta indica* | Nimbiol |
| Fusarium wilt | *Fusarium oxysporum* | Avr1 | Virulence effector | *Azadirachta indica* | 6-Deacetylnimbin |
| Fusarium wilt | *Fusarium oxysporum* | Avr1 | Host–pathogen interaction | *Azadirachta indica* | Gedunin |
| Fusarium wilt | *Fusarium oxysporum* | Avr1 | Effector-mediated pathogenicity | *Azadirachta indica* | Sugiol |
| Fusarium wilt | *Fusarium oxysporum* | Avr1 | Disease establishment | *Azadirachta indica* | Kulinone |

**Table 2.** Molecular docking results of phytochemicals against Avr1 protein.

| Rank | Ligand ID | Binding Affinity (kcal/mol) | RMSD (Upper Bound, Å) | RMSD (Lower Bound, Å) |
| --- | --- | --- | --- | --- |
| 1 | ECN2_IMPHY000093 | -6.3 | 0.00 | 0.00 |
| 2 | ECN2_IMPHY000093 | -6.2 | 16.676 | 13.993 |
| 3 | ECN2_IMPHY000093 | -6.0 | 5.753 | 3.743 |
| 4 | ECN2_IMPHY000093 | -6.0 | 29.907 | 26.777 |
| 5 | ECN2_IMPHY000093 | -5.9 | 6.363 | 3.936 |
| 6 | ECN2_IMPHY000093 | -5.6 | 23.891 | 22.351 |
| 7 | ECN2_IMPHY000093 | -5.6 | 5.461 | 2.508 |
| 8 | ECN2_IMPHY000093 | -5.6 | 5.702 | 2.975 |
| 9 | ECN2_IMPHY000093 | -5.5 | 5.702 | 2.985 |
| 10 | ECN2_IMPHY000093 | -5.5 | 15.577 | 13.617 |

### 3.2 ADME Analysis

Pharmacokinetic profiling was conducted to assess the drug-likeness and bioavailability of the top ten compounds. ADME analysis indicated that nimbiol and gedunin exhibited favorable pharmacokinetic properties, including high gastrointestinal absorption and compliance with Lipinski’s rule of five (Table 3). Several other compounds, such as 6-deacetylnimbin and kulonate derivatives, showed moderate absorption, while compounds like kulactone and kulolactone displayed lower predicted absorption. These results highlight nimbiol and gedunin as potential lead compounds with suitable drug-like characteristics for further investigation.

**Table 3.** ADME and drug-likeness properties of top-ranked compounds.

| Compound | MW (g/mol) | HB A | HB D | iLOG P | Log S | GI Absorption | BB B | Lipinski i | PAINS |
| --- | --- | --- | --- | --- | --- | --- | --- | --- | --- |
| Nimbiol | 272.38 | 2 | 1 | 2.86 | -5.22 | High | Yes | Yes | 0 |
| 6-Deacetylnimbin | 498.56 | 8 | 1 | 3.78 | -4.84 | High | No | Yes | 0 |
| Gedunin | 482.57 | 7 | 0 | 3.23 | -5.75 | High | No | Yes | 0 |
| Sugiol | 300.44 | 2 | 1 | 3.29 | -5.65 | High | Yes | Yes | 0 |
| Methyl 2,5-dihydroxycinnamate | 194.18 | 4 | 2 | 1.59 | -1.42 | High | Yes | Yes | 0 |

### 3.3 Toxicity Prediction

The top ten ligands were evaluated for predicted toxicity parameters, including hepatotoxicity, carcinogenicity, immunotoxicity, mutagenicity, and cytotoxicity. Most compounds demonstrated moderate to high immunotoxicity, with nimbiol and gedunin exhibiting high immunotoxicity scores (0.98 and 0.99, respectively) but acceptable hepatotoxicity and mutagenicity profiles (Table 4). These findings suggest that, while the compounds may require careful consideration for potential immunotoxic effects, they generally possess a favorable toxicity profile suitable for further exploration as antifungal agents.

**Table 4.**
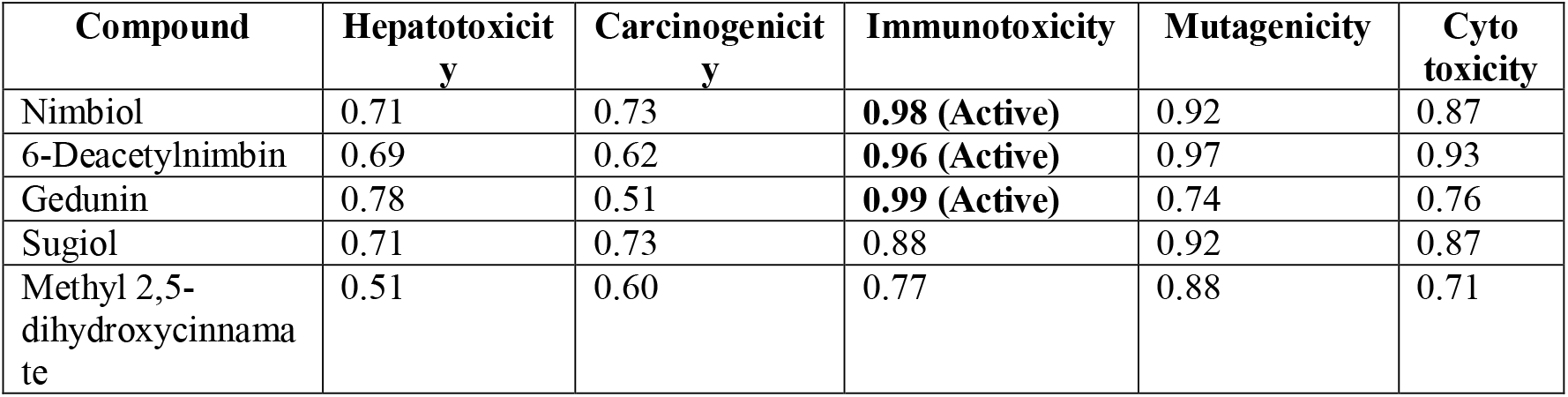
Toxicity prediction profile of selected phytochemicals.

| Compound | Hepatotoxicity | Carcinogenicity | Immunotoxicity | Mutagenicity | Cyto toxicity |
| --- | --- | --- | --- | --- | --- |
| Nimbiol | 0.71 | 0.73 | **0.98 (Active)** | 0.92 | 0.87 |
| 6-Deacetylnimbin | 0.69 | 0.62 | **0.96 (Active)** | 0.97 | 0.93 |
| Gedunin | 0.78 | 0.51 | **0.99 (Active)** | 0.74 | 0.76 |
| Sugiol | 0.71 | 0.73 | 0.88 | 0.92 | 0.87 |
| Methyl 2,5-dihydroxycinnamate | 0.51 | 0.60 | 0.77 | 0.88 | 0.71 |

### 3.4 Protein–Ligand Interaction Analysis

The highest-ranking protein–ligand complex, Avr1–nimbiol, was further analyzed to identify the key interactions stabilizing the complex. Multiple hydrophobic and electrostatic interactions were observed with residues ARG180, VAL158, LYS182, and TYR134 (Table 5). ARG180 formed a strong electrostatic Pi-cation interaction, while other residues contributed hydrophobic contacts, including Pi-alkyl and alkyl interactions. These interactions indicate a stable binding mode and suggest that nimbiol could potentially interfere with Avr1 effector activity.

**Table 5.** Key interactions of the highest-ranking Avr1–ligand complex.

| Protein Residue | Ligand Atom | Distance (Å) | Interaction Category | Interaction Type |
| --- | --- | --- | --- | --- |
| ARG180 | UNK1 | 4.52 | Electrostatic | π-Cation |
| ARG180 | UNK1 | 5.33 | Hydrophobic | Alkyl |
| VAL158 | UNK1 | 4.28 | Hydrophobic | Alkyl |
| LYS182 | UNK1 | 4.11 | Hydrophobic | Alkyl |
| TYR134 | UNK1 | 5.44 | Hydrophobic | π-Alkyl |
| TYR134 | UNK1 | 5.16 | Hydrophobic | π-Alkyl |

### 3.5 Summary of Results

In summary, nimbiol and gedunin demonstrated the highest binding affinities and favourable pharmacokinetic profiles among the ten screened neem-derived compounds. Toxicity predictions indicated moderate hepatotoxicity and high immunotoxicity, warranting further experimental validation. Interaction mapping revealed stable hydrophobic and electrostatic contacts with key Avr1 residues, supporting the potential inhibitory effect of nimbiol against Fusarium effector activity. These findings provide a strong rationale for the further development of these compounds as lead candidates for managing Fusarium wilt.

**Figure 01.**
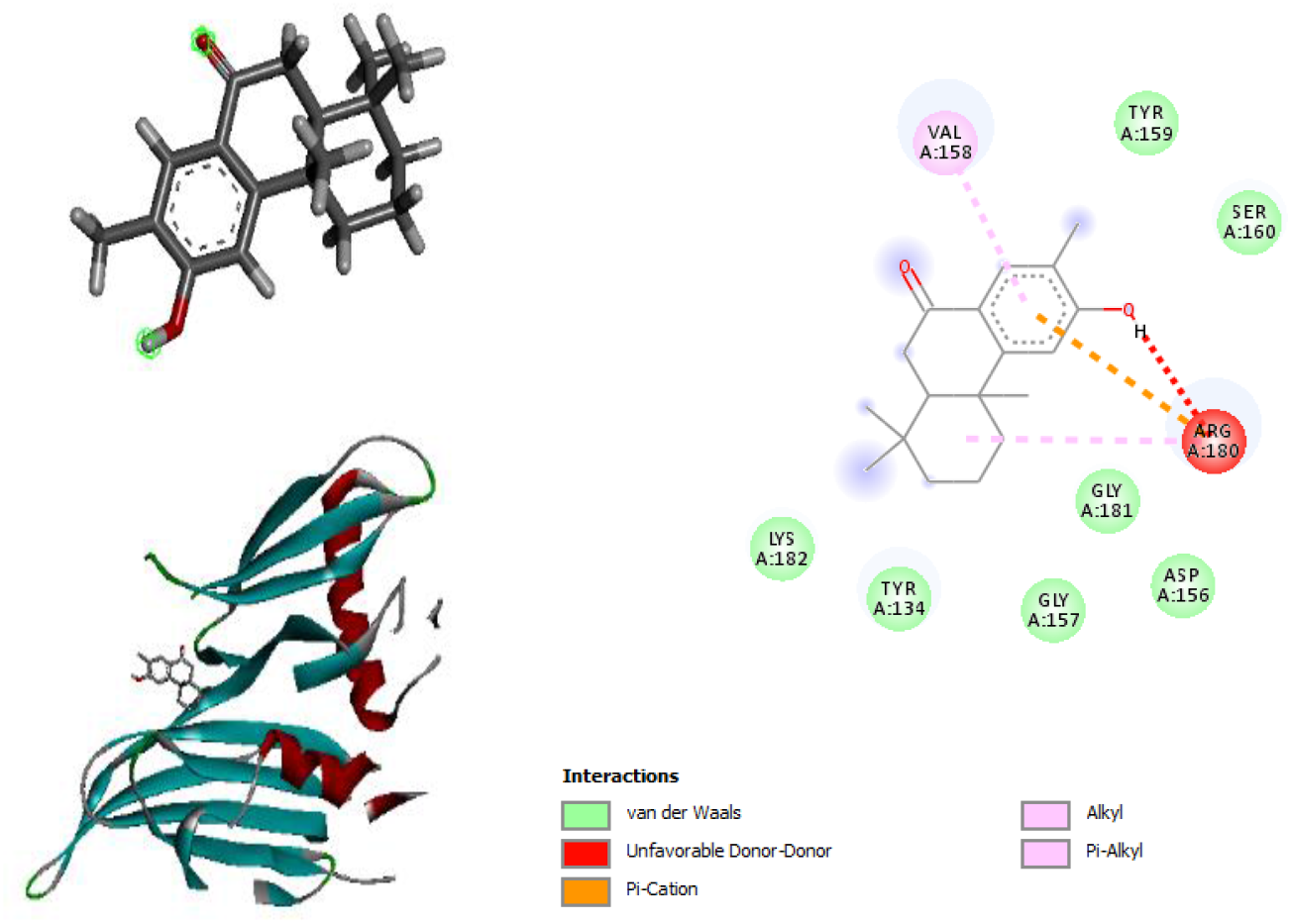
Three-dimensional binding conformation of the top-ranked neem-derived compound within the active site of the Avr1 effector protein, highlighting key interacting residues.

## 4.0 Discussion

The present study investigated the binding affinities, pharmacokinetic properties, toxicity profiles, and molecular interactions of neem-derived phytochemicals against the Avr1 (SIX4) effector protein of *Fusarium oxysporum*, aiming to identify potential lead compounds for managing Fusarium wilt. Our integrative in silico approach identified nimbiol and gedunin as top candidates with favorable profiles.

### 4.1 Antifungal Potential of Neem Phytochemicals

Previous studies have reported that *Azadirachta indica* (neem) extracts possess significant antifungal activity against *F. oxysporum* (Janja, 2025; Bhattacharyya & Jha, 2011; Fernandes et al., 2019; Abdelgaleil et al., 2005). In vitro analyses demonstrated that neem leaf extracts significantly inhibited the mycelial growth of *F. oxysporum*, suggesting that neem-derived compounds can effectively suppress pathogen development under laboratory conditions (N.O. Agbenin & Marley, 2006). For example, neem extract showed up to ∼83% inhibition of *F. oxysporum* mycelial growth compared to control treatments in agar assays, confirming its antifungal potency among various medicinal plant extracts (Dissanayake, 2014).

Neem’s antifungal efficacy has also been associated with its rich content of bioactive limonoids (e.g., azadirachtin, nimbin, gedunin), which have been shown to interfere with fungal cell wall synthesis and spore germination. Previous observations (Shubham et al., 2025; Braga et al., 2020; Houterman et al., 2008)) provide biological support for our docking results, where nimbiol and gedunin displayed the highest binding affinities toward Avr1, suggesting that these compounds may contribute to the inhibitory effects reported in empirical studies.

### 4.2 Comparison with Other Bioactive Targets

While most studies involving neem’s antifungal activity focus on crude extracts or in vitro growth inhibition (Shubham et al., 2025; Mahmoud et al., 2011), limited research has directly targeted specific pathogen virulence proteins. A recent molecular docking study investigated Bacillus-derived compounds against several *F. oxysporum* virulence factors, including Avr1 (SIX4), and found strong binding affinities, indicating that targeting effector proteins is a promising bioactive strategy (K. Vigneshwaran et al., 2025).

Our results align with this emerging trend: nimbiol and gedunin showed affinities in the range expected for biologically relevant interactions, supporting the concept of targeting effector proteins as part of antifungal strategies. This integrative in silico focus goes beyond conventional antifungal activity assays by providing a mechanistic hypothesis for how neem phytochemicals may modulate effector function.

### 4.3 ADME and Drug-Likeness Profiles

Pharmacokinetic prediction revealed that nimbiol and gedunin possess high gastrointestinal absorption and satisfy Lipinski’s rule of five without PAINS alerts, suggesting potential bioavailability and reduced risk of nonspecific interference. These findings are consistent with previous in silico and empirical reports of plant limonoids exhibiting favorable pharmacokinetic behaviors and biological activities. Although literature specifically reporting ADME data for these compounds against plant targets is sparse, the general drug-likeness of limonoids has been described in phytochemical research due to their balanced physicochemical properties (Shubham et al., 2025; Alsaleh et al., 2024; Vélez et al., 2022; Dwivedi et al., 2021).

### 4.4 Toxicity Profiles and Safety Considerations

Toxicity predictions demonstrated moderate hepatotoxicity but generally favorable profiles for nimbiol and gedunin, with no significant mutagenicity or cytotoxicity predicted. High immunotoxicity scores observed across most compounds indicate the need for cautious interpretation, as computational toxicity models can overestimate certain endpoints. However, when taken in the context of known low toxicity of neem applications in agricultural use, these results suggest that *in vivo* studies would be important next steps to clarify safety profiles.

### 4.5 Protein–Ligand Interaction Mechanisms

Detailed interaction mapping of the top-ranking Avr1–nimbiol complex revealed key hydrophobic and electrostatic interactions involving residues such as ARG180, VAL158, LYS182, and TYR134. These interactions suggest stable binding within the effector’s putative active site and are consistent with similar docking studies where fungal virulence proteins formed networks of hydrophobic contacts that underpinned ligand stability (K. Vigneshwaran et al., 2025).

### 4.6 Broader Context and Implications

Collectively, the antifungal activity of neem extracts reported in empirical literature and the mechanistic insights from computational docking reinforce the potential of neem phytochemicals as bioactive agents against *F. oxysporum*. While most existing studies focus on crude extracts or general growth inhibition, our study provides molecular-level evidence suggesting that specific neem compounds may target virulence effectors, an advance in mechanistic understanding.

Moreover, recent agricultural reports confirm that neem foliar extracts or botanical formulations can reduce disease incidence and pathogen proliferation in crops afflicted by Fusarium wilt ((Maheswari et al., 2024; Amit Prakash Kapadnis & Rutuja S. Abhonkar, 2022). Besides, another empirical finding also lends field-level relevance to the molecular interactions we observed computationally (Yi et al., 2021).

The alignment of computational predictions with documented antifungal activities of neem extracts supports the hypothesis that nimbiol and gedunin are promising candidates for further investigation. By bridging *in silico* insights with existing empirical evidence, this study advances understanding of neem phytochemicals as potential effectors of Fusarium wilt management.

## 5.0 Conclusion

This study employed an integrated computational approach to identify neem-derived phytochemicals targeting the Avr1 effector protein of *Fusarium oxysporum*, a major causal agent of Fusarium wilt. Among the screened compounds, nimbiol and gedunin exhibited the strongest binding affinities, favorable ADME profiles, and acceptable predicted toxicity, highlighting their potential as lead antifungal candidates. Interaction analysis revealed stable hydrophobic and electrostatic contacts with key Avr1 residues (ARG180, VAL158, LYS182, and TYR134), suggesting a plausible mechanism for effector inhibition. These findings align with previous reports on neem’s antifungal activity while providing molecular-level insights into specific effector targeting. Overall, the results support the potential of neem-derived phytochemicals as sustainable, plant-based agents for Fusarium wilt management. Further experimental validation, including in vitro and in planta studies, is required to confirm their efficacy and optimize their safety and performance.

## 6.0 Declarations

### Consent to Participate

Not applicable.

### Consent to Publish

Not applicable.

### Ethics Approval and Consent to Participate

Not applicable, as this study did not involve human participants, animals, or clinical samples.

### Clinical Trial Registration

Not applicable.

### Data Availability Statement

All data generated or analyzed during this study are included in this published article and its supplementary information. Docking outputs, ADME, and toxicity prediction results are available from the corresponding author upon reasonable request.

